# Characterizing brain imaging features associated with ADAS-Cog13 sub-scores with 3D convolutional neural networks

**DOI:** 10.1101/2022.03.17.484832

**Authors:** Kaida Ning, Pascale B. Cannon, Jiawei Yu, Srinesh Shenoi, Lu Wang, Alzheimer’s Disease Neuroimaging Initiative, Joydeep Sarkar

**Affiliations:** Holmusk Inc., Singapore

**Author notes:** For the Alzheimer’s Disease Neuroimaging Initiative Data used in preparation of this article were obtained from the Alzheimer’s Disease Neuroimaging Initiative (ADNI) database (adni.loni.usc.edu). As such, the investigators within the ADNI contributed to the design and implementation of ADNI and/or provided data but did not participate in analysis or writing of this report. A complete listing of ADNI investigators can be found at: http://adni.loni.usc.edu/wpcontent/uploads/how_to_apply/ADNI_Acknowledgement_List.pdf. **CORRESPONDING AUTHOR** Correspondence to Joydeep Sarkar.

## Abstract

To date, few advanced machine learning models have been developed for investigating the associations between features from brain imaging and individual Alzheimer’s disease (AD) related cognitive functional changes. Additionally, how these associations differ among different imaging modalities is unclear. Here we investigated 3D convolutional neural network (CNN) models which were trained to predict sub-scores in 13-item Alzheimer’s Disease Assessment Scale - Cognitive Subscale (ADAS-Cog13) based on MRI and FDG-PET brain imaging data obtained from the ADNI database. We found that each key ADAS-Cog13 sub-score was associated with a specific set of brain features within an imaging modality. Overall, sub-scores were strongly associated with structural changes of subcortical regions including amygdala, hippocampus, and putamen, and were associated with metabolic changes of cortical regions including the cingulated gyrus, occipital cortex, middle front gyrus, precuneus cortex, and the cerebellum. Our findings provided insights into complex AD etiology. Our analytical pipeline can also be utilized to study other brain diseases.

## INTRODUCTION

Alzheimer’s Disease (AD) is the most frequent cause of dementia^1^. AD pathology is characterized by the accumulation of toxic species, such as amyloid beta plaques and tau tangles, alterations in glucose metabolism, as well as brain atrophy ^2^. The progression of AD impacts an individual’s cognitive functions such as memory, language, and spatial navigation ^3,4^. The pathological changes of AD brain is best captured through neuroimaging techniques, such as magnetic resonance imaging (MRI) for brain structural changes, positron emission tomography (PET) for metabolic and chemical composition changes ^5,6^, etc. On the other hand, the cognitive function of AD patients can be evaluated via Alzheimer’s Disease Assessment Scale-Cognitive subscale (ADAS-Cog), which quantifies cognitive functions from different aspects (e.g., word recall, orientation, comprehension, etc.) with continuous values, and is frequently used in research and clinical settings^7,8^.

Study of brain imaging data is important for understanding AD etiology and improving AD diagnosis, prognosis^9,10^, and development of treatments. To date, researchers have identified brain imaging biomarkers that are strongly associated with AD diagnosis: atrophy in the hippocampus and the medial temporal lobe, hypometabolism of glucose in the cingulate cortex, etc. ^10–13^. Further, advanced statistical models have been trained to accurately classify AD versus healthy control brains based on imaging data ^14,15^. However, most studies focused on AD diagnosis instead of individual components of cognitive function, which assesses brain functions from various aspects (e.g., as quantified by ADAS-Cog13 sub-scores). Associations between individual cognitive functions and brain imaging features remain to be further studied. Further, how these associations appear in different imaging modalities such as MRI vs PET remain to be studied. To the best of our knowledge, no previous study has systematically investigated these questions.

To understand the relationship between ADAS-Cog13 sub-scores and brain imaging features, we first linked these two types of data through a statistical model. Here we chose to use a 3-dimensional (3D) convolutional neural network (CNN) model to predict ADAS-Cog13 sub-scores, since CNNs demonstrated to have superior performance in classification and regression when dealing with imaging data ^16,17^. We trained the model, validated it, and further investigated the model. While neural networks (NNs) were commonly used as “black-box” tools in the past, recent advances in methods for interpreting NNs allow researchers to identify features important for the models’ performance ^18–20^. In this study, we applied occlusion ^18–20^, a commonly used method, to investigating the trained model and identified brain features most important for predicting ADAS-Cog13 sub-scores.

In this study, we obtained MRI, fluorodeoxyglucose PET (FDG-PET), and AV45-PET imaging data, along with ADAS-Cog13 sub-scores for subjects from Alzheimer’s Disease Neuroimaging Initiative (ADNI) database (http://adni.loni.usc.edu). We identified top ADAS-Cog13 sub-scores that were most important for AD diagnosis, then used a same pipeline to train CNN models for predicting these key ADAS-Cog13 sub-scores based on different imaging modalities. We further investigated these trained models to identify brain regions associated with ADAS-Cog13 sub-scores. Our analytical pipeline brought new insights for associations between brain features and individual cognitive functions and can be applied to studying other brain diseases where imaging data are available.

## RESULTS

### Identifying ADAS-Cog13 sub-scores important for AD diagnosis

We identified the ADAS-Cog13 sub-scores most important for AD diagnosis by training a random forest (RF) model for classifying AD versus non-AD (nAD) patients based on these sub-scores using ADNI sample (See Methods section and Supplementary Table 1). The RF classifier was able to classify the AD diagnosis with an accuracy of 95% (98% precision and 90% recall). In this model, top four ADAS-Cog13 sub-scores were responsible for 82% of the RF feature importance: word recall (Q1), 16%; delayed word recall (Q4), 29%; orientation (Q7), 25%; word recognition (Q8), 11% (Supplementary Table 2). These top four sub-scores scores are representable of the AD cognitive performance and are therefore used as regression outputs of the subsequent 3D CNN models.

### Predicting key ADAS-Cog13 sub-scores with CNNs based on brain images

We trained 3D CNNs that utilized brain images to predict 4 ADAS-Cog13 sub-scores (Q1, Q4, Q7, and Q8) based on MRI, FDG-PET, and AV45-PET imaging modalities. Our CNN model architecture is described in Figure 1 (see Methods section for details). We found that the MRI based CNN model performed the best in predicting sub-scores, with R^2^ of 78%, 80%, 64%, and 62% for Q1, Q4, Q7 and Q8 respectively (Figure 2D). The FDG-PET based 3D CNN performed similarly to the MRI model, with the exception of poor performance on Q8 (49%). The AV45-PET based model had the lowest prediction accuracy (Figure 2D).

**Figure 1.**
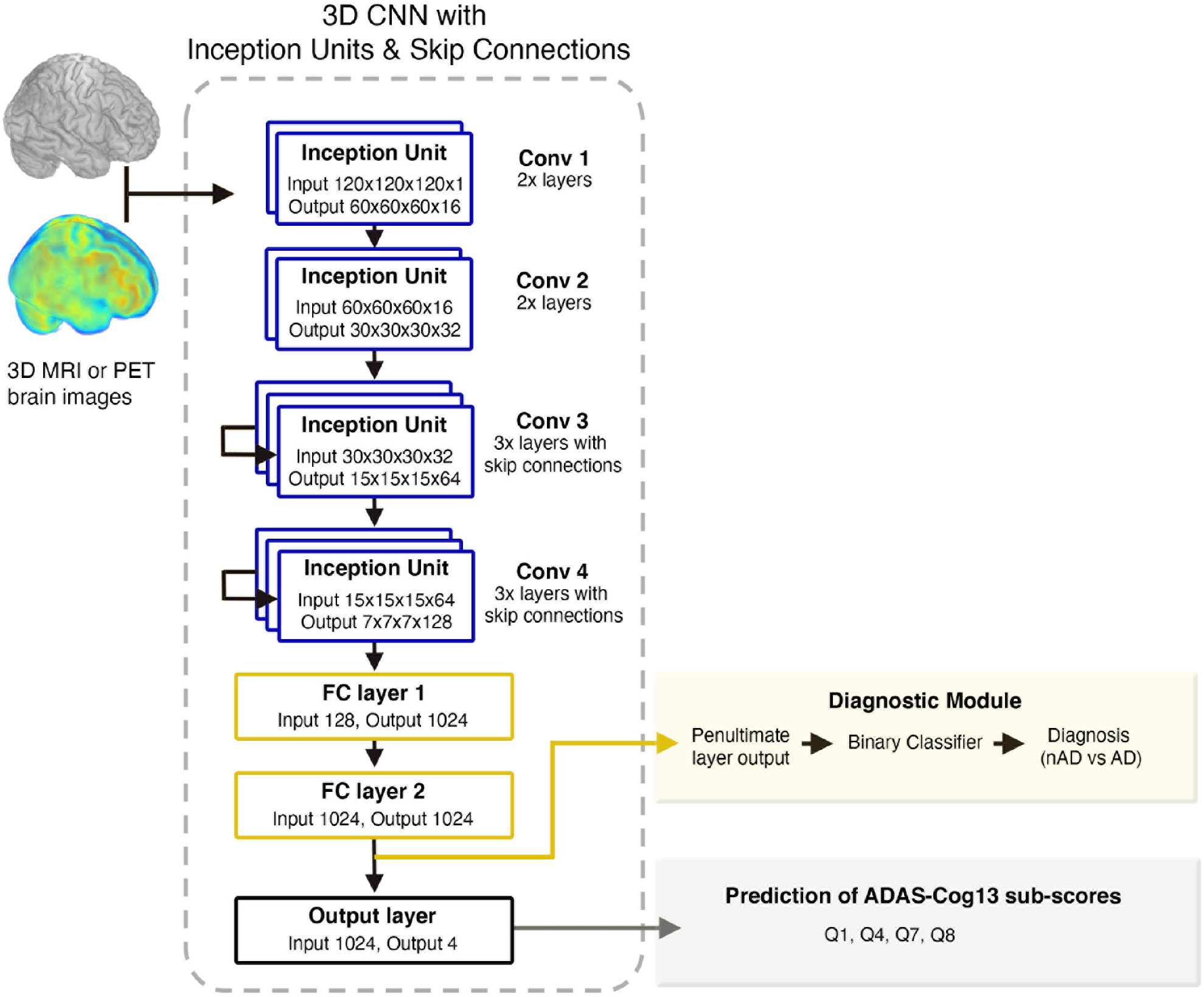
3D CNN model architecture. The CNN model is composed of 10 convolutional layers grouped: Conv 1 (2 layers), Conv 2 (2 layers), Conv 3 (3 layers) and Conv 4 (3 layers). This model processes brain imaging data, and can predict both AD diagnosis (yellow box) and ADAS-Cog13 sub-scores (gray box).

**Figure 2.**
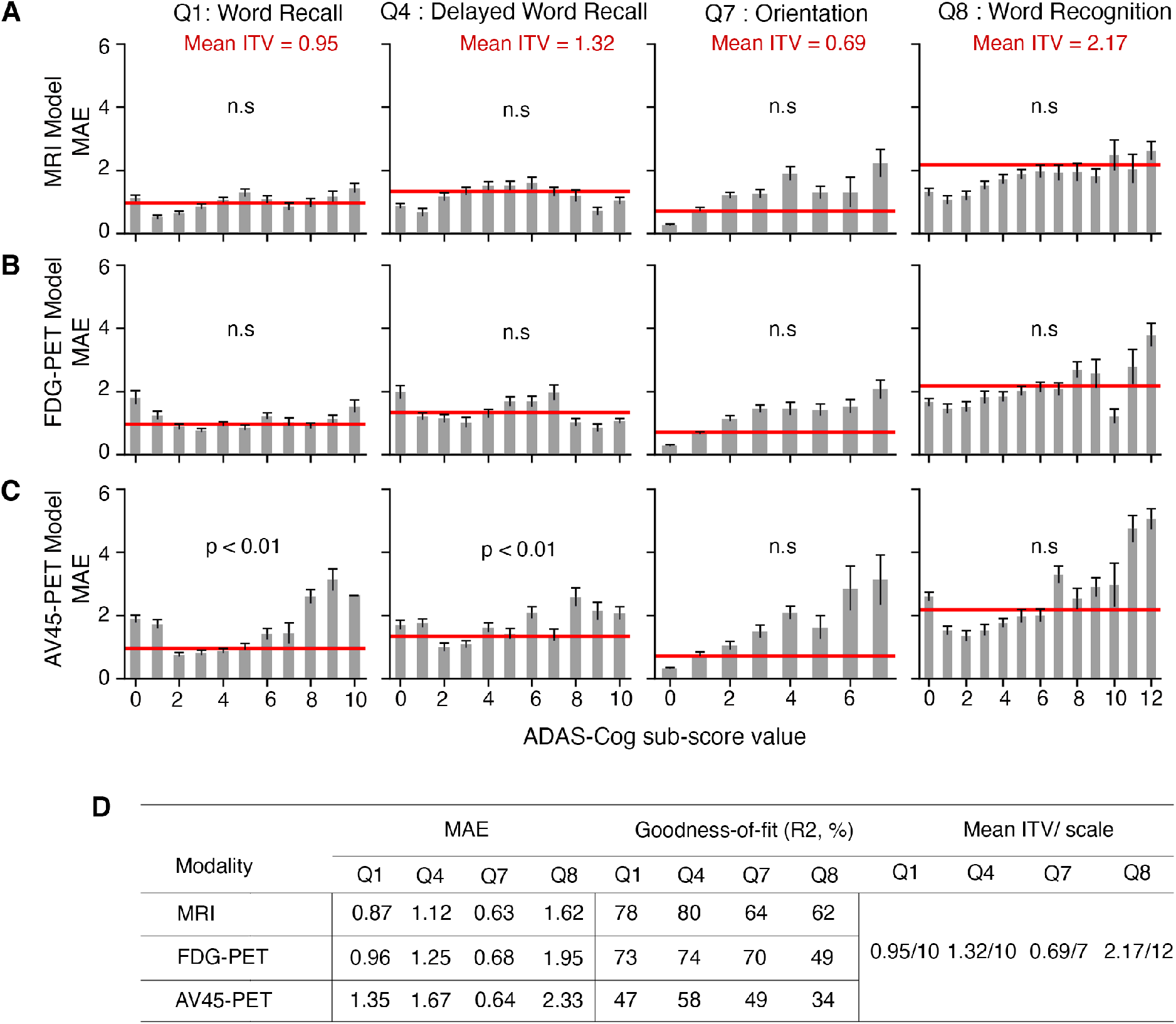
Model metrics on ADAS-Cog13 predictions. A-C: Graphical representation of the mean absolute error (MAE) across ADAS-Cog sub scores for the MRI (A), FDG-PET (B) and AV45-PET (C) CNNs. The inter-test variabilities (ITVs) for four ADAS-Cog13 sub-scores are represented by the red horizontal line. One-tailed t-tests was performed to compare MAEs and ITVs for each ADAS-Cog13 sub-score. ‘n.s.’ means there is no significant difference between MAE and ITV. D: Test set MAE and R^2^ for the MRI, FDG-PET and AV45-PET based CNN models, along with the mean ITV and scale of each ADAS-Cog13 sub-score.

To assess the models’ accuracy in predicting ADAS-Cog13 sub-scores, we compared the mean absolute error (MAE) of our model predictions with the inter-test variability of sub-scores (ITV), which reflects natural fluctuations of ADAS-Cog13 (see Methods section for details). The MAEs of MRI and FDG based 3D CNN model didn’t show significant difference from ITVs for Q1, Q4, Q7, or Q8 (Figure 2 A, B, D). This indicated that errors in the model predictions are comparable to intrinsic variations in the sub-scores. As a comparison, AV45-PET based model performed worse, with MAEs significantly higher than ITVs for Q1 and Q4 (Figure 2 C, D).

To further assess the 3D CNN models’ accuracy, we extended them for classifying nAD vs. AD (i.e., AD diagnosis). The deep learning features from the final fully connected layer (FC2) were used for AD classification (Figure 1). The highest classification accuracy was achieved by the FDG-PET model with a k-nearest neighbor (KNN) extension (AUROC = 90%), followed by the MRI model (AUROC = 89%) and AV45-PET model (AUROC = 84%) (Supplementary Figure 1, Supplementary Table 3). To further test the generalizability of the 3D CNN models, an external validation was done by applying our models to the RADC dataset (see Methods section and Supplementary Table 4) to classify nAD vs. AD. Despite significant differences in age, overall patient populations and scanning protocols between ADNI and RADC datasets, our model achieved an AUROC of 0.74 for RADC data.

### Identifying brain regions associated with ADAS-Cog13 sub-scores

After model training, we investigated MRI and FDG-PET based models with occlusion method to identify brain regions important for predicting ADAS-Cog13 sub-scores (see Methods section for details).

We found that the MRI and FDG-PET CNNs utilized different brain regions for predicting ADAS-Cog13 sub-scores. Further, each ADAS-Cog13 sub-score was associated with a specific set of brain features. In the MRI based 3D CNN model, sub-score Q1 was most strongly associated with brain structural changes in the hippocampus and the putamen, etc. Q4, Q7, and Q8 were strongly associated with changes in the hippocampus and the amygdala, etc. (Figure 3 A-D). Three regions (the amygdala, the hippocampus, and the putamen) appeared among the top ten for all four ADAS-Cog13 sub-score. Supplementary Table 5 lists feature importance scores of all brain regions in the MRI based CNN. In the FDG-PET based 3D CNN model, Q1 and Q4 were most strongly associated with brain metabolic changes in the cerebellum and the cingulate gyrus (posterior division), etc. Q7 was strongly associated with changes in the cingulate gyrus (posterior division) and the thalamus, etc. Q8 was associated with changes in the cingulate gyrus (posterior division) and the putamen, etc. (Figure 3 E-H). Five brain regions appeared among the top ten for all four ADAS-Cog13 sub-scores: the cingulate gyrus (posterior division), middle frontal gyrus, precuneous cortex, lateral occipital cortex (inferior division), and cerebellum. Supplementary Table 6 lists feature importance scores of all brain regions in the FDG-PET based CNN. Figure 3 I-L visualize the top five regions associated with each ADAS-Cog13 sub-score in MRI model (red) and FDG-PET model (green), respectively. A few important brain regions overlap between MRI and FDG-PET based CNNs, as highlighted in yellow color.

**Figure 3.**
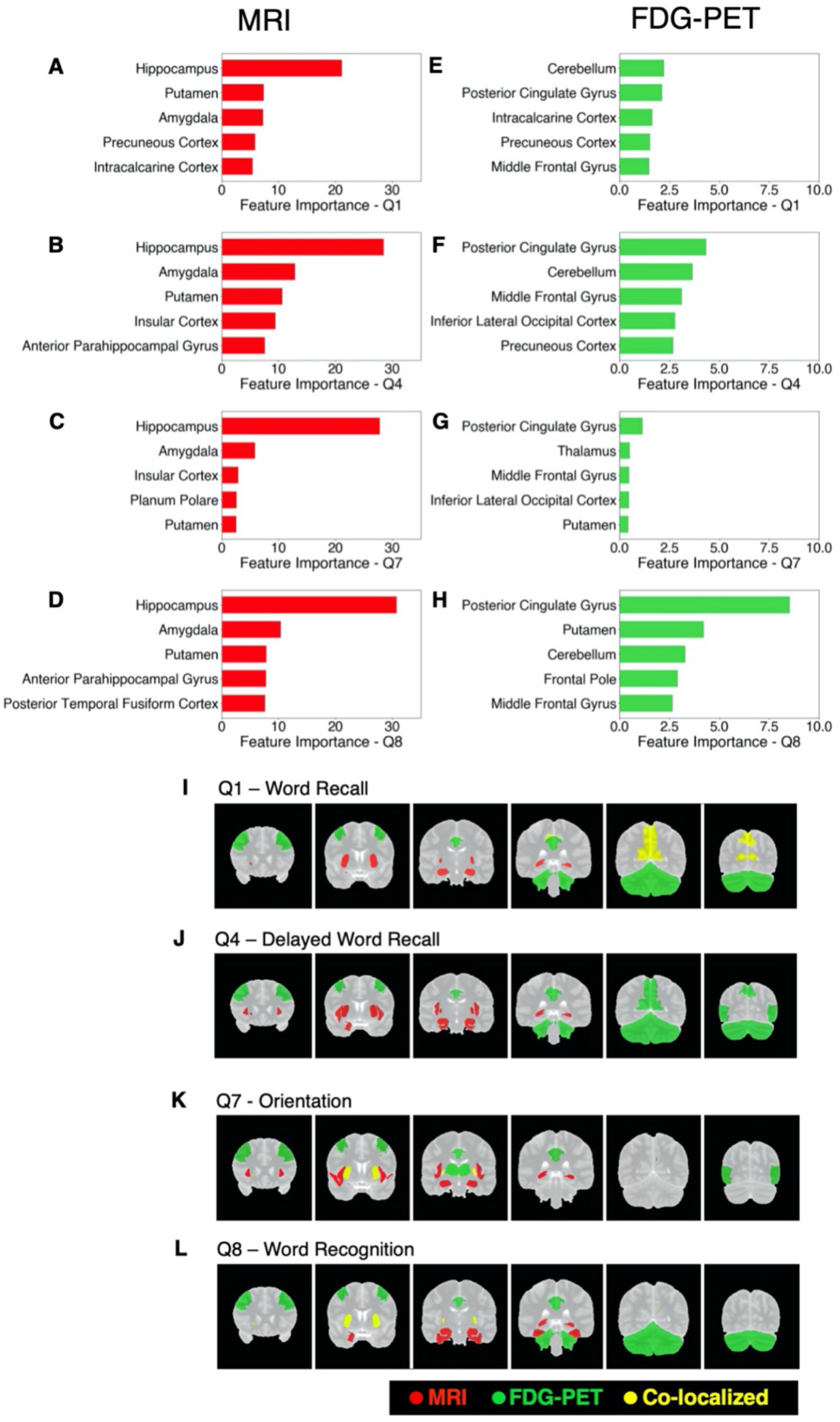
Important brain regions in MRI and FDG-PET based 3D CNN models. A-D: Feature importance scores of top five brain regions for ADAS-cog13 sub-scores Q1, Q4, Q7, and Q8 in the MRI based CNN. E-H: Feature importance scores of top five brain regions for Q1, Q4, Q7, and Q8 in FDG-PET based CNN. I-L: Coronal views of top five important brain regions for each ADAS-Cog13 sub-scores in the MRI (red) and the FDG-PET (green) 3D CNNs. Regions that are important for both MRI and FDG-PET models are highlighted in yellow colour.

We further investigated the brain-imaging associations in three disease sub-groups: cognitively normal (CN), MCI, and AD. We observed that within different disease sub-groups, the importance scores of a specific brain region varied. For example, the hippocampus was a most important region in the MRI based CNN model in CN subjects for all ADAS-Cog13 sub-scores, while its importance in the model diminished in MCI and AD patients. The cerebellum was a most important region in the FDG-PET model in AD subjects for sub-score Q7; while its importance was much lower in CN and MCI subjects (Supplementary Figure 2).

### Pairwise correlation among ADAS-Cog13 sub-scores

We also calculated pairwise correlation among ADAS-Cog13 sub-scores Q1, Q4, Q7, and Q8 in terms of their associations with brain features (see Methods section for details). In the MRI based model, the strongest correlation was between Q1 and Q4 (Spearman’s correlation=0.94), followed by Q4 and Q8 (Spearman’s correlation=0.91). In comparison, Q7 had lower correlations with other sub-scores. All pairwise correlations are shown in Figure 4

**Figure 4.**
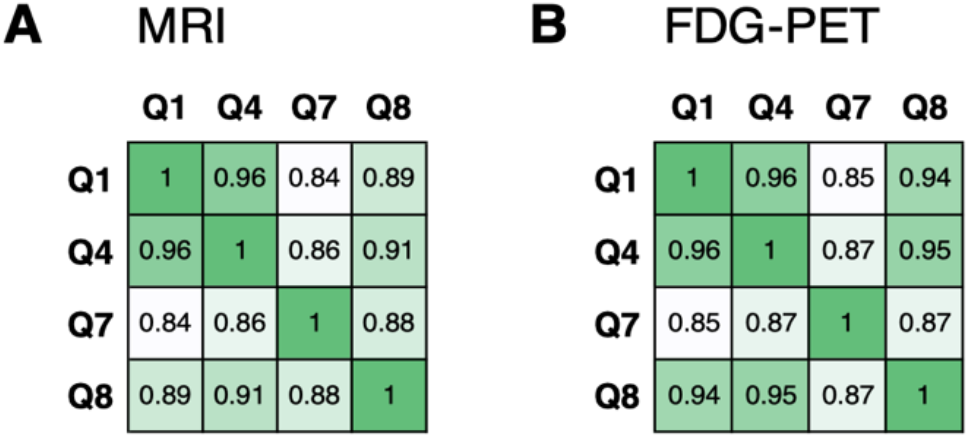
Pairwise correlations among ADAS-Cog13 sub-scores Q1, Q4, Q7, and Q8. A: Correlations in MRI based CNN. B: Correlations in FDG-PET based CNN.

A. Note that Q1 measures function for word recall, Q4 measures delayed word recall, Q7 measures orientation, and Q8 measures word recognition. The pairwise correlations of sub-scores based on brain feature importance scores were higher within language related sub-scores than those between language and orientation sub-scores. In the FDG-PET model, the strongest pairwise correlation of cognitive functions was between Q1 and Q4 (Spearman’s correlation=0.96), followed by Q4 and Q8 (Spearman’s correlation=0.95). All pairwise correlations are shown in Figure 4 B. Like the observation for the MRI based model, the pairwise correlations were higher within language related sub-scores than those between language and orientation related sub-scores.

## DISCUSSION

In this study, we made a first attempt to train 3D CNN models based on brain imaging data for predicting four ADAS-Cog13 sub-scores that were crucial for AD diagnosis, and then investigated the CNN models to identify brain regions strongly associated with these sub-scores.

The MRI and FDG-PET based 3D CNN models predicted the ADAS-Cog13 sub-scores with R^2^ above 60%, except for low performance of FDG-PET based model on predicting sub-score Q8. The MAEs of these models’ prediction on ADAS-Cog13 sub-scores were comparable to the ITVs of these sub-scores, indicating that errors in CNN models’ prediction were comparable to natural variations of the ADAS-Cog13 sub-scores. We provided additional internal validation of the MRI and FDG-PET based models by demonstrating that they can be applied to classifying nAD vs. AD subjects accurately (with an AUROC of 0.89 and 0.9, respectively). Further, the MRI based model performed well in classifying nAD vs. AD subjects when applied to an external test dataset, RADC, without any modifications. Compared to MRI and FDG-PET based CNN models, the AV45-PET CNN model performed worse in predicting ADAS-Cog13 sub-scores but had comparable performance in identify AD vs nAD. As revealed in previous studies, amyloid deposition as measured by AV45-PET has an impact on cognition in early stages, while ADAS-Cog13 is not sensitive enough for measuring changes in early cognitive stage of MCI or AD ^21,22^. This is a potential explanation for poor performance of AV45-PET based model in predicting ADAS-Cog13 sub-scores. In real-world practice, many clinical trials for AD drugs monitor cognitive endpoints like ADAS-Cog13 and amyloid beta based on AV45-PET imaging ^23^. Our observation suggests that cognitive functions had a stronger association with MRI and FDG-PET imaging signals than with AV45-PET imaging signals. Additionally, MRI and/or FDG-PET neuroimaging biomarker monitored during drug treatments can provide valuable information on change in brain structure & metabolism in response to treatment and overall progression of disease.

After training MRI and FDG-PET based 3D CNN models, we identified brain regions associated with key ADAS-Cog13 sub-scores through investigating these models with occlusion method. Thanks to the statistical nature of this method, we were able to quantify the contribution of each brain region on prediction of ADAS-Cog13 sub-scores, which was not encoded in other brain regions. We found that these models utilized distinct sets of brain regions for predicting the sub-scores. For example, the hippocampus region had a high importance score in predicting all ADAS-Cog13 sub-scores in MRI based CNN model (Figure 3, Supplementary Table 5). This is a subcortical region important for memory formation and is well known to undergo atrophy in AD patients ^10,13^. In comparison, the hippocampus region didn’t appear to be highly important for the FDG-PET based CNN model. Instead, a network of cortical regions, led by cingulate gyrus, appeared to be highly important for all the sub-scores in the FDG based model (Supplementary Table 6). This finding corroborated previous studies that reported abnormal metabolism in cingulate cortex of AD patients ^11,12^. Further, we found that cerebellum, which is essential for motor activity and motor learning, was an important region associated with cognitive functions, especially Q1, Q4, and Q8, in the FDG-PET modality. Previous studies have reported that metabolites of cerebellar neurons promote amyloid-β clearance, and that cerebellar glucose metabolism was significantly lower in AD patients compared to control subjects ^24,25^.

Our study also showed that within an imaging modality (MRI or FDG-PET) each ADAS-Cog13 sub-scores were associated with a specific set of brain regions. In the MRI based model, sub-score Q1 was most strongly associated with brain structural changes in the hippocampus and the putamen, etc. Q4, Q7, and Q8 were strongly associated with changes in the hippocampus and the amygdala, etc. In the FDG-PET based model, Q1 and Q4 were most strongly associated with brain metabolic changes in the cerebellum and the cingulate gyrus (posterior division), etc. Q7 was strongly associated with changes in the cingulate gyrus (posterior division) and the thalamus, etc. Q8 was associated with changes in the cingulate gyrus (posterior division) and the putamen, etc. These findings indicate a complex underlying relationship between structural and functional changes in brain regions (as measured by brain biomarkers) and changes in specific cognitive functions as observed in AD etiology. Nevertheless, the cognitive function pairs that were similar to each other were highly correlated in terms of their associations with brain features as well. We further made a first attempt to investigate the CNN models within each disease sub-group and found that ranks of brain region importance scores were different among disease sub-groups (Supplementary Figure 2). For example, the hippocampus, a most important region in the MRI based model in CN subjects for all ADAS-Cog13 sub-scores, showed lower importance score in MCI and AD patients. This indicated that AD etiology is dynamic, with different brain regions becoming strongly associated with cognitive function as the disease progresses.

Our study had some limitations. First, we chose our 3D CNN model structure and parameters based on previous knowledge on training CNNs. CNNs have lots of variations in their structures and parameters. Exploring more combinations of CNN structures and parameters may improve the model’s accuracy in predicting ADAS-Cog13 sub-scores. Second, our definition of brain feature importance was based on occlusion method, while alternative definitions such as GRAD-RAM are available and may reveal other insights ^20,26^. Third, our current model predicted cognitive functions collected at a single time point. A natural extension of this model would be to incorporate time factor, so that it predicts the change of cognitive function in the future.

In clinical practice, our findings may help to refine the process of AD early interventions and clinical trials. It is known that the changes of the brain, although associated with cognitive function changes, can occur a long time before changes in cognitive function. For example, it was reported that brain structural changes were detectable in the hippocampus and the medial temporal lobe up to ten years before any AD symptom arises ^27^. Also, researchers were able to predict progression from mild cognitive impairment to AD two years in advance using FDG-PET or MRI data ^14,28^. Based on our analyses, we further suggest that brain features identified in our model, along with the cognitive scores predicted based on brain imaging data, may assist AD risk assessment before diagnoses, allowing early disease intervention. During patient enrollment for clinical trials, our model may also help to stratify the patients in terms of their disease progression risk and increase the power of these trials. Most such current applications use more traditional radio-imaging features like volume, average grey value etc., which ignore the deeper associations in the grey-values across the 3D space. CNN models can capture deeper associations and generate more nuanced brain feature based patient stratification. We further propose that brain structural and metabolic features be monitored after initiation of drug intervention: changes of these features, while highly associated with ADAS-cog sub-scores, may occur well before any change of cognitive functions and can therefore suggest AD stabilization (or even reversion) and help clinicians to better understand and evaluate drug efficacy.

In sum, we developed 3D CNN models for analyzing 3D brain imaging data. These models predicted ADAS-Cog13 sub-scores based on different imaging modalities and brought insights for the association between cognitive functions and brain imaging features. Our models can hopefully accelerate clinical trials for AD and can be further expanded to analyze imaging data for different types of brain diseases.

## MATERIALS AND METHODS

### ADNI data

The data used was obtained from the Alzheimer’s Disease Neuroimaging Initiative (ADNI) database (http://adni.loni.usc.edu). Data used in the preparation of this article were obtained from the Alzheimer’s Disease Neuroimaging Initiative (ADNI) database (adni.loni.usc.edu). The ADNI was launched in 2003 as a public-private partnership, led by Principal Investigator Michael W. Weiner, MD. The primary goal of ADNI has been to test whether serial magnetic resonance imaging (MRI), positron emission tomography (PET), other biological markers, and clinical and neuropsychological assessment can be combined to measure the progression of mild cognitive impairment (MCI) and early Alzheimer’s disease (AD). For up-to-date information, see www.adni-info.org. As specified by the ADNI protocol, each participant within the study was willing, spoke English or Spanish, was able to perform all test procedures described in the protocol and had a study partner able to provide an independent evaluation of functioning. We used 9,862 unique imaging entries with cognitively normal (CN), mild-cognitive impairment (MCI) or AD diagnosis from ADNI. Demographic and clinical characteristics of these entries are shown in Supplementary Table 1.

### Random forest (RF) classifier for Alzheimer’s Disease (AD) vs non-AD (nAD)

A random forest model^29^ was used to classify AD versus nAD with 13 ADAS-Cog13 sub-scores as predictors. MCI and CN participants were grouped into the nAD class. To balance the weights of the 13 ADAS-Cog13 sub-scores, each sub-score was normalized to a scale of 0 to 1 using min-max scaling. Train / test splitting with a ratio of 80:20 was carried out before model training process. The training and testing sets were balanced by random sampling to adjust for the ratio of nAD and AD. During the training process, model hyperparameters were optimized using grid search ^30^.

### Imaging pre-processing

3D brain images used in this study underwent multiple preprocessing steps as illustrated in Supplementary Figure 3. First, images were registered with the MNI152 standard space structural brain template ^31^. Brain volumes and positions were standardized. Second, a standardized brain mask was applied to strip the cranium and brain stem, retaining only the cerebrum and cerebellum.

For MRI imaging data, white stripe normalization ^32^ was applied. The normal-appearing white matter (NAWM) with least pathological variation was selected as the reference tissue ^32^. All MRI images were then transformed by matching their distributions of NAWM to that of the reference MRI with a fixed mean (*μ*_ref_) and standard deviation (*σ*_ref_), as shown in Supplementary Figure 3. Mathematically, the histogram distribution of a specific MRI image was transformed using

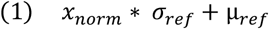

where *x*_*norm*_ = (*x* − μ)/*σ* is the normalized distribution with a mean of zero and a standard deviation of 1 for NAWM. *μ* and *σ* are the mean and standard deviation of NAWM from the MRI image to be normalized. Note that some MRI images failed white stripe normalization, showing a mismatched NAWM peak following normalization (Supplementary Figure 3). Abnormal and low-quality MRI images were excluded from subsequent experiments.

For PET imaging data, normalization was done following the ADNI protocol, where the intensities were pre-normalized by the ratio between the radiotracer and the body weight ^33^. In addition, we applied a customized “cohort normalization” to scale PET images into a range between 0 to 1 at a cohort level. Mathematically, all PET images were divided by the maximum of *n*_stats_ from the training cohort, where *n*_stats_ was the voxel intensity at the 99.9 percentile from the PET image from an individual patient. We used 99.9 percentile instead of the maximum intensity value was to avoid the influence from outliers.

### Building 3D convolutional neural networks (3D CNNs)

We built a 3D convolutional neural network (3D CNN) with skip connection and inception units to analyze 3D brain imaging data. The backbone of the 3D-CNN is based on VGG16 ^34^, which consists of 14 convolutional (conv) layers (including 4 max pooling layers), followed by 2 fully connected layers (FC1 and FC2) and a final output layer which generates predictions of four ADAS-Cog13 sub-scores as in a multi-task feature learning process ^35^. When training the model, a batch size of 8 imaging samples was used. Rectified linear unit (ReLU) was applied as the activation functions for all the conv layers. Batch normalization (BN) was applied before the activation function. The channel number (*n*_channel_) for the conv layers inside Conv 1, Conv 2, Conv 3 and Conv 4 were 16, 32, 64 and 128. At the end of Conv 1, Conv 2, Conv 3 and Conv 4, a max pooling operation with kernel size of 2 were applied to reduce the spatial size of activation maps into half. Conv 3 and Conv 4 were modified into residual blocks ^36^ with skip connections which aggregate the output of the 1^st^ and 2^nd^ conv layers in the group before feeding the results into the 3^rd^ conv layer. The output of the Conv 4 with a size of (*n*_batch_, 7, 7, 7, 128) was flattened into an array with size (*n*_batch_, 43,904) before proceeding to the fully connected (FC) layers: FC1 and FC2. Both FC1 and FC2 contained 1024 neurons which converted the input array into arrays of size (*n*_batch_, 1024). The tanh activation function was applied to both FC layers. Finally, the output layer with a customized sigmoid activation function (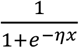 where *η* was a trainable parameter) that converts the FC2 layer output into the final predictions of size (*n*_batch_, 4).

To train the model, the samples were split into training and test sets with a ratio of 80:20. The mean squared error (MSE) between the true ADAS-Cog13 scores and the predicted scores were used as the loss function. Model parameters were then optimized using the Adam algorithm ^37^ to minimize the MSE of the training set. Cosine annealing ^38^ was used as the learning rate scheduler to help the model converge rapidly to a local minima and at the same time prevent the model from getting stuck in one single local minima by abruptly increasing the learning rate to maximum at the beginning of each cycle. The maximum and minimum learning rate used in our study were 0.01 and 0.0001, respectively. To boost the model accuracy, ensemble technique was applied by selecting 5-7 best models (models with minimum MSE on the test set) from the saved models and calculated the mean of predictions from multiple models for each ADAS-Cog13 sub-score.

To evaluate model performance, we used multiple metrics including the mean absolute error (MAE) and R^2^. The MAE for each ADAS-Cod sub-score was defined as the mean of |*y*_true_-*y*_pred_| across the cohort where *y*_true_ and *y*_pred_ are the true and predicted ADAS-Cog13 scores.

To compare the model performance with clinical practice, we calculated the inter-test variability (ITV) for each ADAS-Cog13 sub-score on a complete ADNI ADAS-Cog13 dataset with 9,862 unique samples (the same cohort as listed in Supplementary Table 1). ITV was calculated as the maximum difference in recorded score for each ADNI participant within a period of +/- 3 months. Mathematically, the ITV for entry *i* (*ITV*_*i*_) was defined as

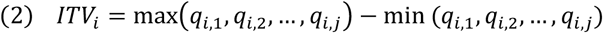

where *q*_*i*,1_, *q*_*i*,2_, …, *q*_*i,j*_ were the ADAS-Cog13 cog sub-score (Q1, Q4, Q7 or Q8) from all the visits that were within +/- 3 months with regards to entry *i*. The mean ITV for each sub-score represents the mean of all ADNI participants’ ITV.

### Diagnostic extension & validation of 3D CNN models

The diagnostic extension of the 3D CNN predicted AD vs. nAD, where the penultimate layer (FC2) output with a size of (*n*_entries_, 1024) was utilized to train a binary classification model, where *n*_entries_ was the number of entries. Classification models, including logistic regression^39^, k-nearest neighbor (KNN) ^40^ and random forest ^29^, were trained to predict diagnosis using the values in the penultimate layer. Similar to the main AI model, multiple sub-models with the same architecture were ensembled and a voting classifier was applied to generate final predictions.

The validation dataset for nAD and AD diagnosis was obtained from the Rush Alzheimer’s Disease Center (RADC) Religious Order Study (ROS) ^41^, a clinical-pathologic study of aging and dementia. The demographic information of samples from RADC is summarized in Supplementary Table 4. MRI images from RADC were processed in the same way as for ADNI images.

### Identifying features important for 3D CNN model

We estimated feature importance scores of 56 brain regions using occlusion method as described in previous studies ^18–20^. The feature importance of a specific brain region quantifies its importance that was not encoded in other brain regions. In our analyses, the cortical and subcortical atlases were obtained from the Center for Morphometric Analysis (https://cma.mgh.harvard.edu). With the occlusion method, the change of model prediction error after removing a brain region was measured. More specifically, the CNN feature importance score of brain region *i* was defined as

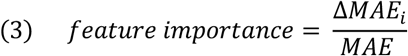

where Δ*MAE*_*i*_ is the absolute change of CNN MAE after removing the *i*^th^ brain region from model input.

### Calculating pairwise correlation among ADAS-Cog13 sub-scores

We calculated pairwise correlation among ADAS-Cog13 sub-scores Q1, Q4, Q7, and Q8 based on their associations with 56 brain regions. To be specific, for each ADAS-Cog13 sub-score we obtained feature importance scores of 56 brain regions in a CNN model. We then calculated Spearman’s correlation between each pair of sub-scores based on the 56-dimentional feature importance scores. The pair-wise correlations among sub-scores were obtained for MRI and FDG-PET based CNN models separately.

## ACKNOWLEDGEMENTS

Data collection and sharing for this project was funded by the Alzheimer’s Disease Neuroimaging Initiative (ADNI) (National Institutes of Health Grant U01 AG024904) and DOD ADNI (Department of Defense award number W81XWH-12-2-0012). ADNI is funded by the National Institute on Aging, the National Institute of Biomedical Imaging and Bioengineering, and through generous contributions from the following: AbbVie, Alzheimer’s Association; Alzheimer’s Drug Discovery Foundation; Araclon Biotech; BioClinica, Inc.; Biogen; Bristol-Myers Squibb Company; CereSpir, Inc.; Cogstate; Eisai Inc.; Elan Pharmaceuticals, Inc.; Eli Lilly and Company; EuroImmun; F. Hoffmann-La Roche Ltd and its affiliated company Genentech, Inc.; Fujirebio; GE Healthcare; IXICO Ltd.;Janssen Alzheimer Immunotherapy Research & Development, LLC.; Johnson & Johnson Pharmaceutical Research & Development LLC.; Lumosity; Lundbeck; Merck & Co., Inc.;Meso Scale Diagnostics, LLC.; NeuroRx Research; Neurotrack Technologies; Novartis Pharmaceuticals Corporation; Pfizer Inc.; Piramal Imaging; Servier; Takeda Pharmaceutical Company; and Transition Therapeutics. The Canadian Institutes of Health Research is providing funds to support ADNI clinical sites in Canada. Private sector contributions are facilitated by the Foundation for the National Institutes of Health (www.fnih.org). The grantee organization is the Northern California Institute for Research and Education, and the study is coordinated by the Alzheimer’s Therapeutic Research Institute at the University of Southern California. ADNI data are disseminated by the Laboratory for Neuro Imaging at the University of Southern California.

## AUTHOR CONTRIBUTIONS

Research conception and design: K.N., P.B.C.

Data analysis and interpretation: K.N., J.Y., S.S, L.W

Manuscript writing: All authors.

Project supervision: J.S.

## COMPETING INTERESTS

The authors declare no competing interests.

## Supplementary Figures and Tables

**Supplementary Figure 1.**
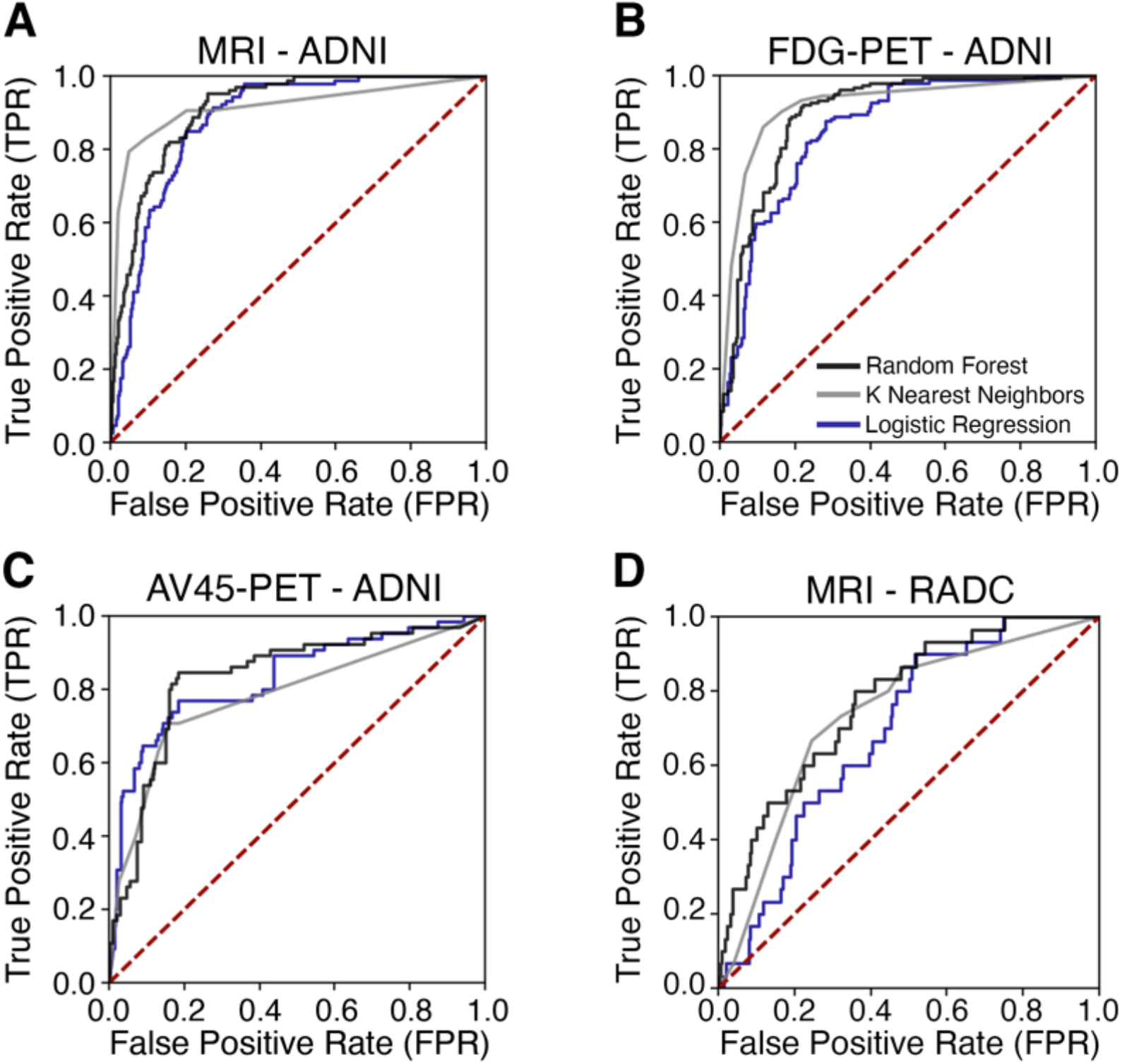
Receiver Operating Characteristic (ROC) curves of classification models on the ADNI and RADC datasets. A-C: MRI, FDG-PET, and AV45-PET based CNN models with diagnostic extension applied to ADNI samples. D: MRI based CNN models with diagnostic extension applied to RADC samples. Random forest (black), K nearest neighbours (grey), and logistic regression (blue) were used for the diagnosis extension.

**Supplementary Figure 2.**
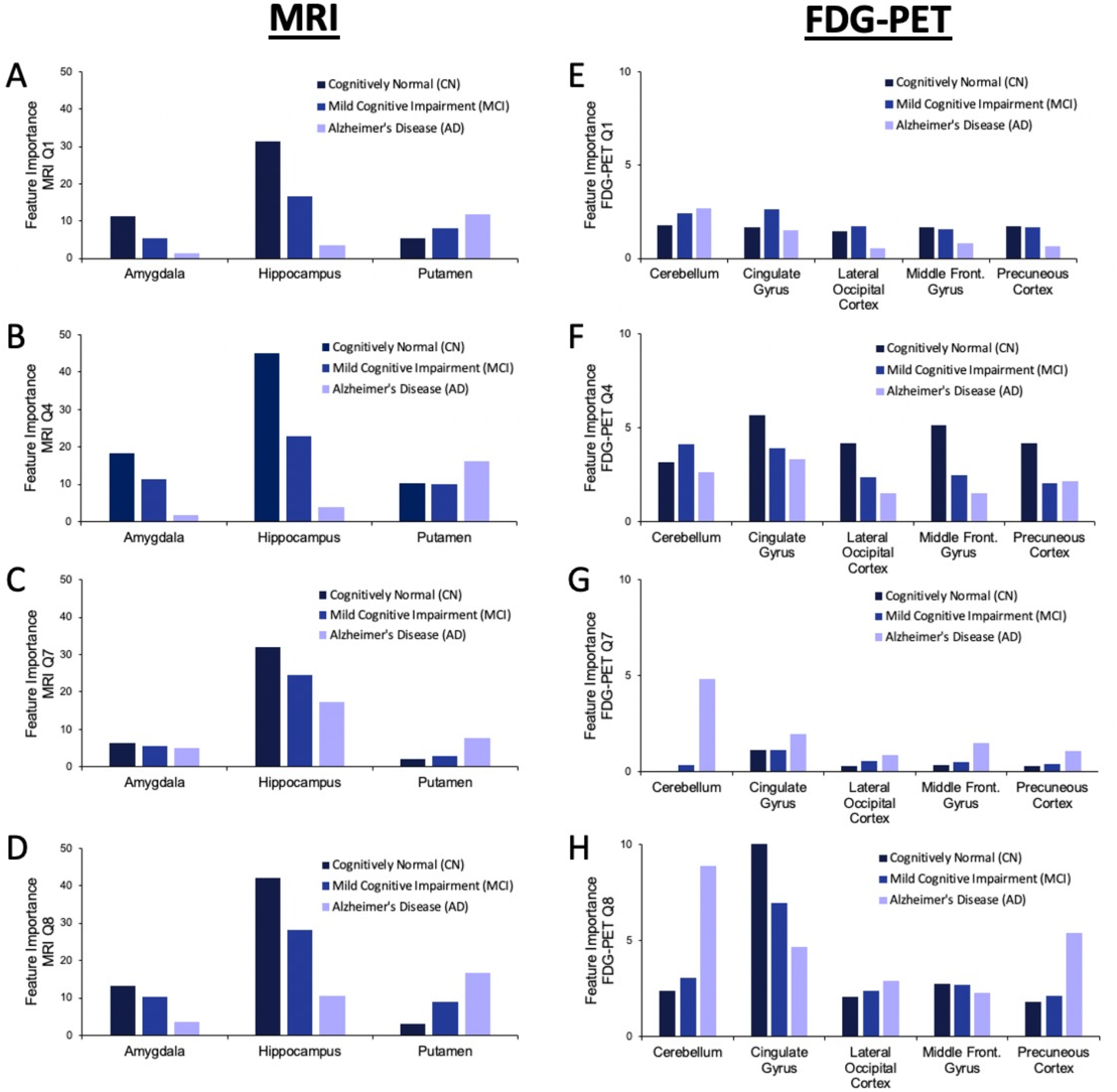
Feature importance scores of selected brain regions within MRI (A-D) and FDG-PET (E-H) in different diagnosis sub-groups.

**Supplementary Figure 3.**
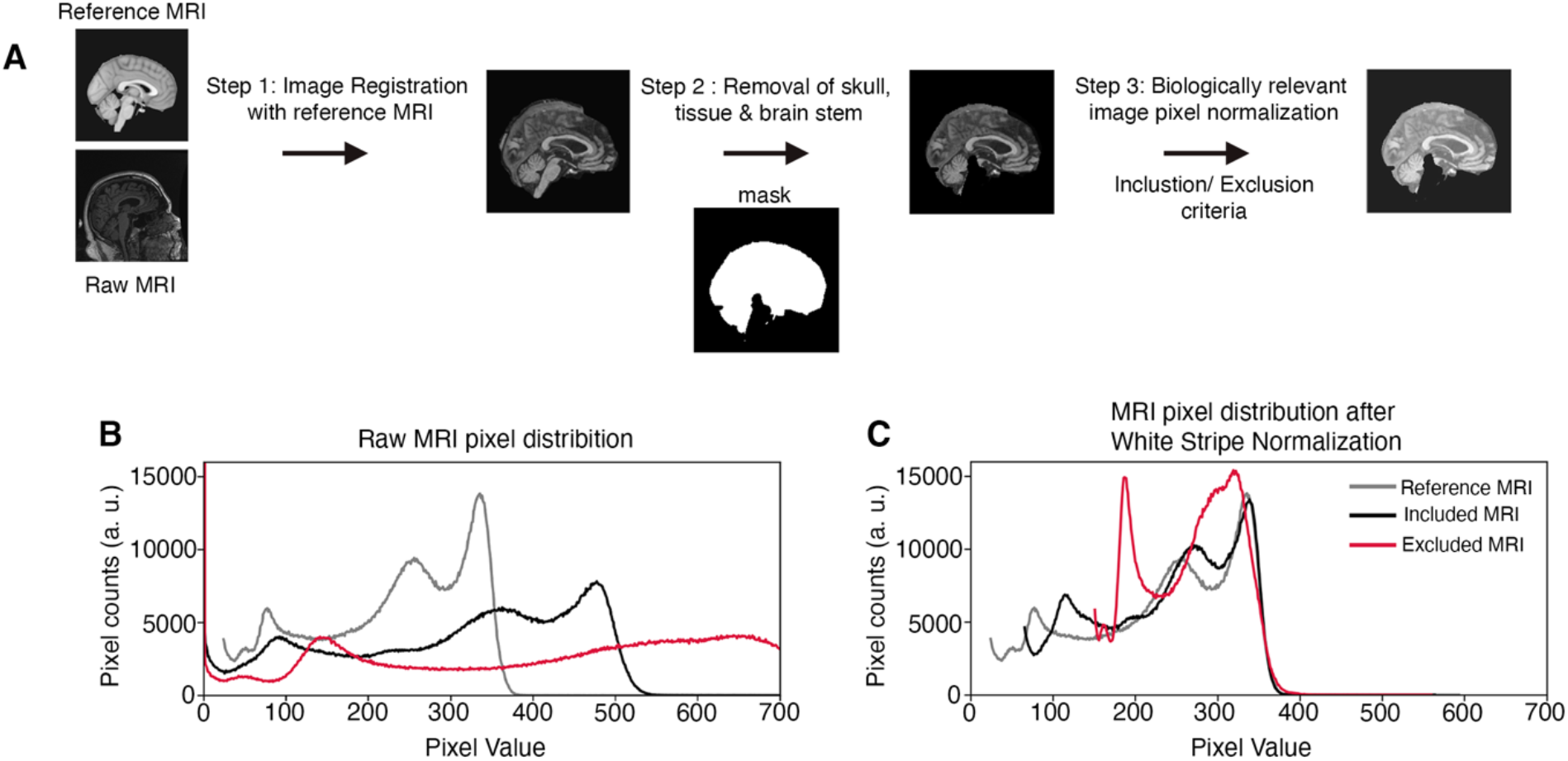
Imaging preprocessing. A) Schematic representation of the image processing pipeline. Example of white-stripe normalization for MRI images, with pixel intensity distributions of the reference MRI (grey lines), a successfully normalized MRI (black lines) and an abnormal MRI that fails white stripe normalization (red lines) from raw imaging (B) to normalized imaging (C).

**Supplementary Table 1.** Demographic information of ADNI samples.

|  |  | **Normal cognition** | **Mild cognitive impairment** | **Alzheimer's Disease** |
| --- | --- | --- | --- | --- |
|  |  | (n = 3314) | (n = 4412) | (n = 2136) |
|  |  | **Count (%)** | **Count (%)** | **Count (%)** |
| **Sex** | **F** | 1722 (52) | 1747 (39.6) | 909 (42.6) |
|  | **M** | **1592(48)** | **2665(60.4)** | **1227(57.4)** |
|  |  | **Mean (stdev)** | **Mean (stdev)** | **Mean (stdev)** |
| **Age** |  | 76.9 (6.6) | 75.1 (7.8) | 76.7 (7.5) |
| **Total ADAS-Cog13** |  | 9.4 (4.6) | 16.3 (7.5) | 32.5 (11.2) |

**Supplementary Table 2.** Random forest feature importance for predicting AD vs. nAD based on ADAS-Cog13 sub-scores.

| **ADAS-COG SUB-SCORE** | **FEATURE IMPORTANCE (%)** |
| --- | --- |
| **Q1 – WORD RECALL** | 16 |
| **Q2 – COMMANDS** | 1 |
| **Q3 – CONSTRUCTIONAL PRAXIS** | 1 |
| **Q4 – DELAYED WORD RECALL** | 29 |
| **Q5 – NAMING OBJECTS/FINGERS** | 2 |
| **Q6 – IDEATIONAL PRAXIS** | 3 |
| **Q7 – ORIENTATION** | 25 |
| **Q8 – WORD RECOGNITION** | 11 |
| **Q9 – REMEMBERING TEST INSTRUCTIONS** | 1 |
| **Q10 – COMPREHENSION** | 1 |
| **Q11 – WORD FINDING LANGUAGE ABILITY** | 2 |
| **Q12 – SPOKEN LANGUAGE ABILITY** | 1 |
| **Q13 – NUMBER CANCELLATION** | 5 |

**Supplementary Table 3.**
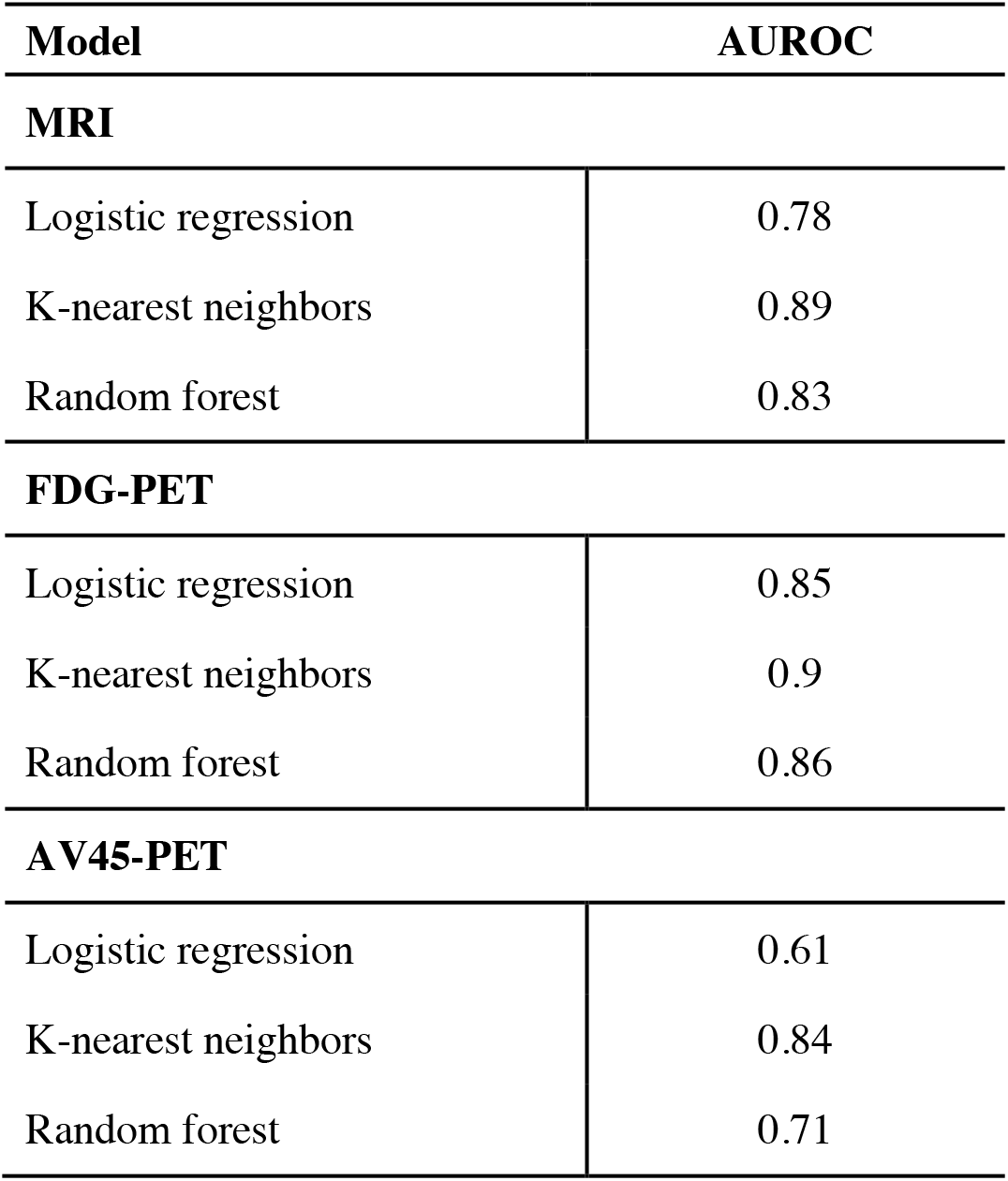
Accuracy of CNN model extension for AD diagnosis.

| **Model** |  | **AUROC** |
| --- | --- | --- |
| **MRI** |  |  |
| Logistic regression |  | 0.78 |
| K-nearest neighbors |  | 0.89 |
| Random forest |  | 0.83 |
| **FDG-PET** |  |  |
| Logistic regression |  | 0.85 |
| K-nearest neighbors |  | 0.9 |
| Random forest |  | 0.86 |
| **AV45-PET** |  |  |
| Logistic regression |  | 0.61 |
| K-nearest neighbors |  | 0.84 |
| Random forest |  | 0.71 |

**Supplementary Table 4.** Demographic information of RADC samples.

|  |  | **Normal cognition** | **Mild cognitive impairment** | **Alzheimer's Disease** |
| --- | --- | --- | --- | --- |
|  |  | (n = 1641) | (n = 325) | (n = 30) |
|  |  | **Count (%)** | **Count (%)** | **Count (%)** |
| **Sex** | **F** | 1271 (77) | 246 (75) | 20 (66) |
|  | **M** | 370 (23) | 79 (24) | 28 (34) |
|  |  | **Mean (stdev)** | **Mean (stdev)** | **Mean (stdev)** |
| **Age** |  | 81.0 (7.3) | 85.0 (6.2) | 86.7 (7.6) |
| **Total MMSE** |  | 28.6 (1.3) | 26.9 (2.1) | 23.0 (3.1) |

## Notes

### Competing Interest Statement

The authors have declared no competing interest.

